# Genetic and Epigenetic Characterization of Sarcoma Stem Cells Across Subtypes Identifies EZH2 as a Therapeutic Target

**DOI:** 10.1101/2024.05.14.594060

**Authors:** Edmond O’Donnell, Maria Muñoz, Ryan Davis, R. Lor Randall, Clifford Tepper, Janai Carr-Ascher

## Abstract

High-grade complex karyotype soft tissue sarcomas (STS) are a heterogeneous and aggressive set of cancers that share a common treatment strategy. Disease progression and failure to respond to anthracycline based chemotherapy, standard first-line treatment, is associated with poor patient outcomes. To address this, we investigated the contribution of STS cancer stem cells (STS-CSCs) to doxorubicin resistance. We identified a positive correlation between CSC abundance and doxorubicin IC_50_ in resistant cell lines. We investigated if a common genetic signature across STS-CSCs could be targeted. Utilizing patient derived samples from five sarcoma subtypes we identified Enhancer of Zeste homolog 2 (EZH2), a member of the polycomb repressive complex 2 (PRC2) responsible for H3K27 methylation as being enriched in the CSC population. EZH2 activity and a shared epigenetic profile was observed across subtypes. Targeting of EZH2 using Tazemetostat, an FDA approved inhibitor specifically ablated the STS-CSC population. Treatment of doxorubicin resistant cell lines with tazemetostat resulted in a decrease in the STS-CSC population. Further, co-treatment was not only synergistic in the parent cell lines, but restored chemosensitivity in doxorubicin resistant lines. These data confirm the presence of shared genetic programs across distinct subtypes of CSC-STS that can be therapeutically targeted.

## Introduction

High-grade complex karyotype sarcomas are treated with aggressive multimodality treatment including surgery, radiation, and chemotherapy. Despite this, the prognosis for these patients is poor with a 65% five-year survival rate. The development of new treatments is complicated by the heterogeneity of disease, evidenced by the more than 70 sarcoma subtypes with varying histology, genetics, and patient demographics^1^. Sarcomas are classified into two genetic groups, simple or complex karyotype. Simple karyotype sarcomas have a known translocation such as EWS-FLI1 that drives Ewing sarcoma^2^. Complex karyotype sarcomas display an aneuploidy phenotype with numerous chromosomal gains and losses^3^. In adults, over 85% of sarcomas have a complex karyotype including undifferentiated pleomorphic sarcoma (UPS), the most common sarcoma subtype. These tumors contain a vast number of chromosomal changes; however, there is a lack of common critical regulatory pathways underlying this aggressive disease^4,5^. Identifying new therapeutic targets across subtypes will result in improved patient outcomes.

Sarcoma mortality is primarily driven by treatment failure and development of metastatic disease. Cancer stem cells (CSCs) are a subpopulation of cells within the bulk tumor that can initiate tumor formation and grow indefinitely^6^. This population has stem cell like properties including protection from apoptosis resulting in therapy resistance and increased metastatic potential^6^. Therefore, there is considerable interest in identifying therapeutic vulnerabilities in sarcoma CSCs as this population is contributing to patient mortality. The majority of patients with high risk localized disease or metastatic sarcoma are treated with doxorubicin, an anthracycline based chemotherapy. Resistance to doxorubicin in sarcoma has been attributed to upregulation of multidrug resistance (MDR) efflux pumps and increased expression of anti-apoptotic factors such as BCL-2^7,8^. In osteosarcoma, resistance to chemotherapy has been attributed to the CSC population and histone deacetylase (HDAC) inhibitors, epigenetic targeting drugs have been shown to sensitize cells to treatment^9,10^. Sarcoma CSCs have been identified across several subtypes such as rhabdomyosarcoma and malignant peripheral nerve sheath tumor (MPNST)^11,12^. Within these sarcomas, CSCs have upregulation of hedgehog, notch, and hippo signaling. In rhabdomyosarcoma, a childhood skeletal muscle tumor, YAP has been shown to promote and maintain stem cells along with NOTCH and SOX^13,14^. An evaluation of CSCs across common complex karyotype sarcomas to identify shared pathways and opportunities for therapeutic targeting has not been performed.

In this study, we observed that doxorubicin resistance in soft tissue sarcomas is associated with an increase in CSC abundance. We hypothesized that a common genetic signature could be identified across CSCs and targeted to overcome resistance. We isolated and genetically evaluated sarcoma CSCs from five common sarcoma subtypes and identified EZH2, a member of the PRC2 epigenetic complex as being upregulated. Targeting of EZH2 led to a reduction of the CSC population and sensitization to doxorubicin indicating this as a potential therapeutic strategy for an aggressive and treatment resistant disease.

## Results

### Acquired resistance to anthracycline chemotherapy correlates with an increase in cancer stem cells

Doxorubicin, an anthracycline-based chemotherapy is the cornerstone of systemic treatment across high-grade complex karyotype sarcoma subtypes. While this is effective, acquired resistance can occur resulting in recurrent or metastatic disease. Resistance to anthracycline therapy is associated with poor prognosis^15^. Across tumor types literature suggests that therapeutic resistance and metastasis are attributable to the cancer stem cell population. To investigate this in sarcoma, we created doxorubicin resistant cell lines. We cultured GCT, a cell line representing undifferentiated pleomorphic sarcoma (UPS) and SW872 derived from dedifferentiated liposarcoma (LPS) with increasing concentrations of doxorubicin (0.05-0.4 uM) over the course of four months. Histologically, xenograft tumors formed from these cell lines were indistinguishable from the parental line (Figure 1A,E). Resistance to doxorubicin (Doxo^R^) was verified with dose-response curves. The doxorubicin resistant GCT (GCT-Doxo^R^) cells had a significantly higher IC50 value as compared to wiltype cells (Figure 1B, 1I). This was also observed in doxorubicin resistant SW872 (SW872-Doxo^R^) cells as compared to WT SW872 lines (Figure 1F, 1I). Importantly, the resistance to doxorubicin was stable in culture over several months in the absence of treatment.

**Figure 1.**
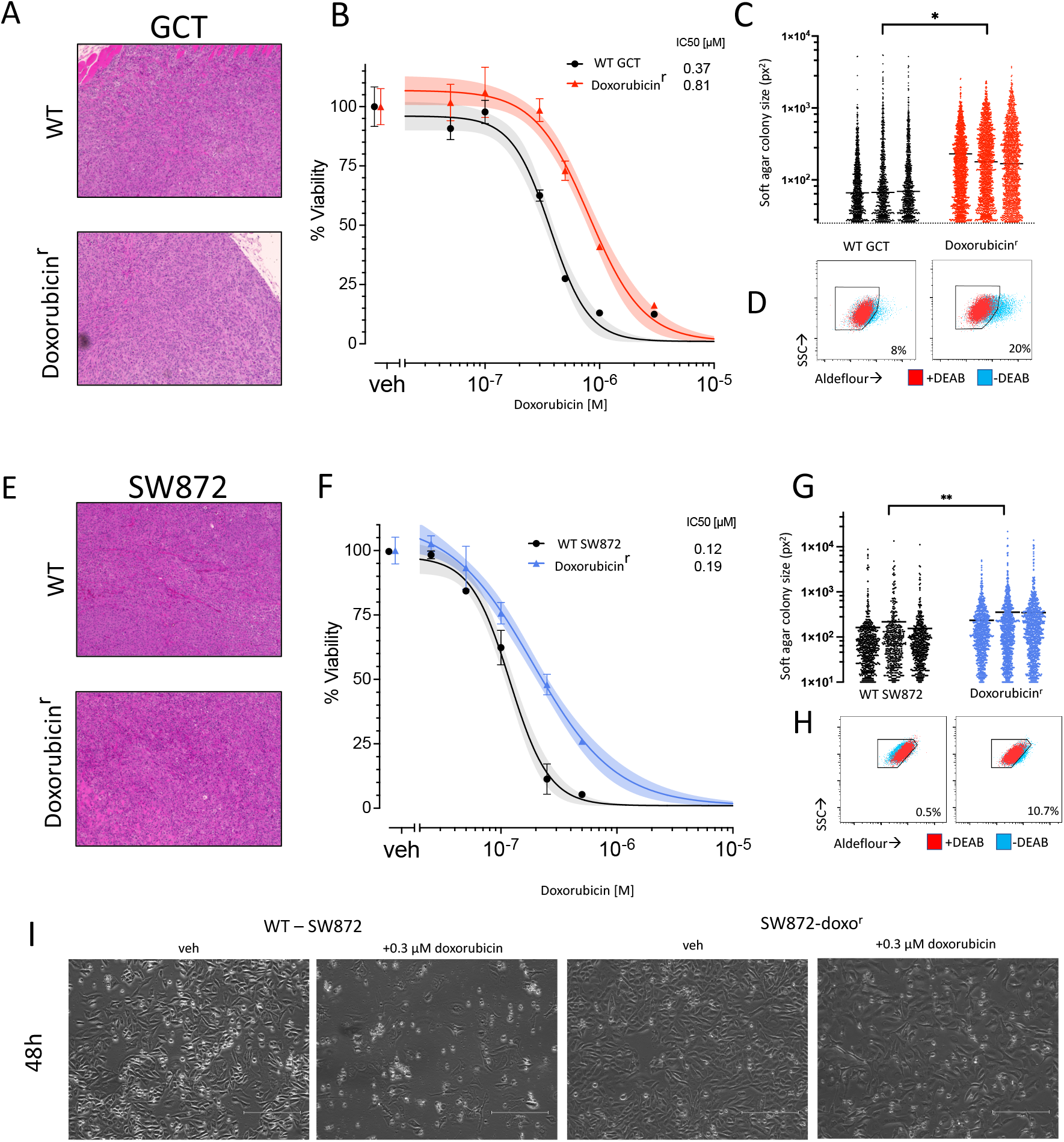
Acquired doxorubicin resistance correlates with increased soft tissue sarcoma cancer stem cell abundance and phenotype. A doxorubicin-resistant (doxo^R^) derivative line of WT GCT (undifferentiated pleomorphic sarcoma) cells was generated through serial passage in increasing concentrations of doxorubicin. A) H&E stains of representative subcutaneous tumors generated from WT and Doxo^R^ SW872 cells. B) Cell-titer glo viability assays for doxorubicin dose-response in WT and doxoR derivative GCT cells. Data points are the mean ± SD of triplicate wells and are representative of three different experiments. C) WT and Doxo^R^ GCT cells were subjected to soft-colony formation assays and the abundance and size of colonies was counted with imageJ software. Each column is a biological replicate. Colony size is shown in units as pixels (px^2^). D) Aldeflour assay showing increased bright population in doxo^R^ (right) cells vs WT GCT cells (left). The DEAB controls are shown in red (n=3). E) H&E stains of representative subcutaneous tumors generated from WT and Doxo^R^ cells SW872 cells (dedifferentiated liposarcoma). F) A doxorubicin-resistant (doxo^R^) derivative line of WT SW872 cells was generated through serial passage in doxorubicin and exhibited an increased IC_50._ Data points are the mean ± SD of triplicate wells and are representative of three different experiments. G) WT and Doxo^R^ SW872 cells were subjected to soft-colony formation assays and the abundance and size of colonies was counted with imageJ software. Colony size is shown in arbitrary units as pixels^2^. G) Aldeflour assay showing increased bright population in doxo^R^ (right) cells vs WT SW872 cells (left) (n=3). I) representative photographs of SW872 WT and Doxo^R^ cells after 48 hours of treatment with either DMSO (vehicle) or 300 nM doxorubicin. bars = 250 nm . *P <0.05, **P<0.05.

To investigate the relationship between doxorubicin resistance and cancer stem cells, we evaluated functional differences in resistant and wildtype cell lines. We first subjected WT and Doxo^R^ cells to soft agar assay given the ability to grow in anchorage independent conditions is associated with a CSC phenotype. We used image-based analysis to quantify the size and abundance of colonies (Figure 1C,G). In both the GCT and SW872 cell model systems, colonies generated by Doxo^R^ derivatives were both more numerous per assay as well as larger in size. This result suggested a more abundant population of cancer stem cells within Doxo^R^ derivative lines compared to the parent cell line following acquisition of chemoresistance. To further evaluate this possibility, we subjected these cell models to the Aldeflour assay, a well-established technique to measure aldehyde dehydrogenase activity of cells. Consistent with a CSC phenotype, there was an increased abundance of Aldeflour bright cells in the SW872 and GCT Doxo^R^ cells (Figures 1D,H).

### Genetic characterization of cancer stem cells across high-grade complex karyotype sarcomas

Acquired resistance to doxorubicin is associated with increased mortality from high-grade sarcomas and we observed this is associated with enrichment of the CSC population. To identify a common target within the CSC population across a diverse subset of sarcomas, we utilized differential gene expression to compare the CSC population to bulk tumor cells. Within sarcomas, there is considerable subtype heterogeneity and a uniform marker of the CSC population is unknown. Across tumor types, aldehyde dehydrogenase (ALDH) is known to be upregulated within CSCs and this has been shown in sarcoma as well. Therefore, we utilized the aldeflour assay, to isolate CSCs from the non-CSC population using cell sorting. To confirm the difference in phenotype, GCT cells were isolated by sorting bright vs. dim cells in the aldeflour assay, and then subjected to soft-agar formation assays. Aldefluor bright GCT cells exhibited better anchorage-independent growth as compared to the Aldefluor dim non-CSC cells (Figure 2A).

**Figure 2.**
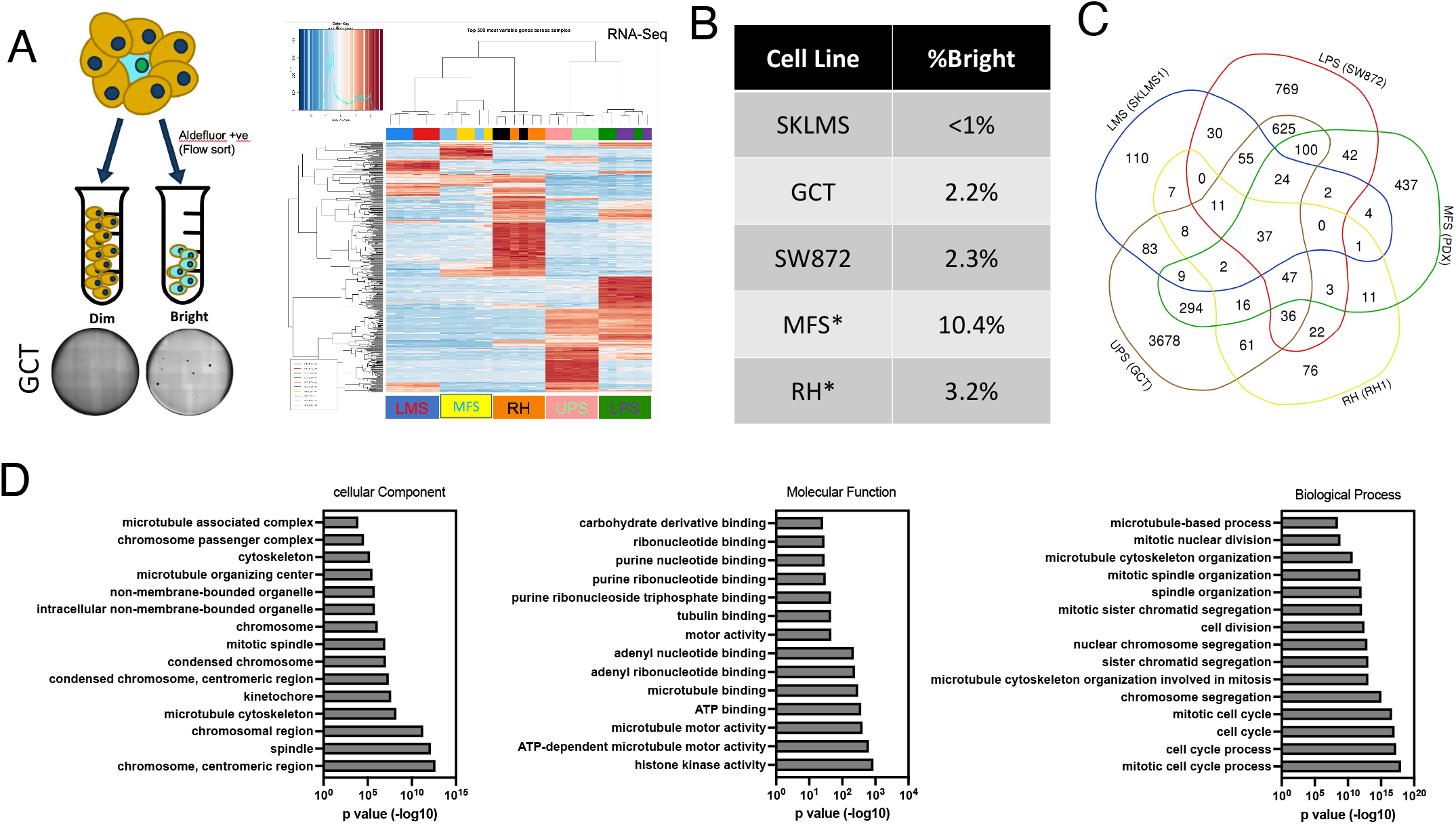
Isolation and RNA-sequencing based analysis of soft-tissue sarcoma cancer stem cells with Aldefluor sorting reveals a shared gene signature. A) Sorting scheme and heatmap of the top 500 differentially regulated genes on RNA-seq analysis of aldeflour bright and dim sorted cells among five soft tissue sarcoma cell lines. A representative soft-agar colony forming assay from sorted GCT cells shows the increased propensity for anchorage-independent growth in aldeflour bright cells. B) Table showing the percent aldeflour positive population among the five different cell lines. LMS=leiomyosarcoma (SKLMS1), RH=rhabdomyosarcoma, MFS=myxofibrosarcoma, UPS=undifferentiated pleomorphic sarcoma (GCT), LPS=dedifferentiated liposarcoma (SW872). B) Percent aldeflour positive (bright) cells for each of the different lines tested. C) Venn diagram showing the overlap of differentially expressed genes with significance p<0.05 (right) and further subdivided according to fold change in expression (left). D) Gene ontology analysis of the 37 shared, differentially regulated genes across the five sarcoma types.

We selected five high-grade complex karyotype sarcoma cell lines that each represent common subtypes. Specifically, leiomyosarcoma, rhabdomyosarcoma, myxofibrosarcoma, undifferentiated pleomorphic sarcoma, and dedifferentiated liposarcoma. Consistent with prior studies, Aldefluor analysis of these cells demonstrated a small number of cells composing the CSC population (Figure 2B)

Aldefluor bright and dim cells from each subtype were sorted and analyzed by RNA-sequencing. Interestingly, differential gene expression showed significant differences when comparing across subtypes but, comparisons of Aldefluor bright to dim cells within sarcoma subtypes demonstrated minimal differences (Figure 2C). To identify differentially regulated genes across bright and dim populations, we utilized a venn-diagram approach of genes significantly upregulated or downregulated at varying fold expression levels. Of note, only one commonly downregulated gene, BRCA, was identified. (Figure 2C). A total of 37 genes were differentially upregulated at an FDR <0.05, comprising 3, 12, and 21 genes by >2, >1.5, and >1.2 fold, respectively. The narrowed gene list (n=37) was evaluated by gene ontology analysis (Figure 2D). We noted significant enrichment for ontology terms related to the maintenance and integrity of cellular mitosis.

### Enhancer of Zeste Homolog 2 (EZH2) is increased in high-grade complex karyotype sarcoma cancer stem cells

To identify potentially targetable pathways in the CSC population as compared to the non-CSC population, we performed gene set enrichment analysis (GSEA) of differentially expressed genes. From this, we identified enhancer of zeste homolog 2 (EZH2), the catalytic subunit of the polycomb repressive complex 2 (PRC2) that facilitates H3K27 methylation as an enriched pathway (Figure 3A).

**Figure 3.**
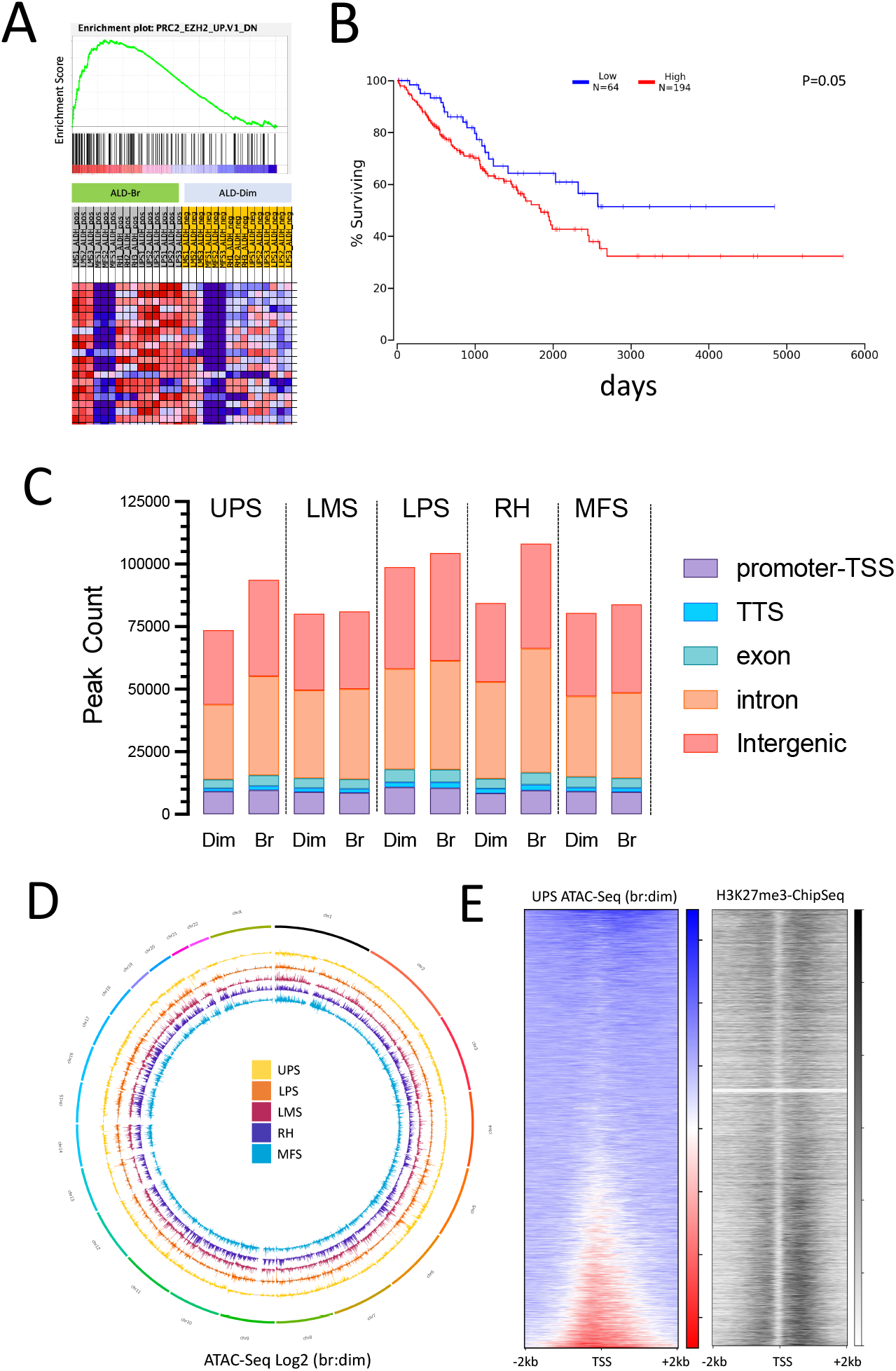
Identification of a shared EZH2 signature among soft tissue sarcoma cancer stem cells. A) Expression data from RNA-Seq was analyzed by gene set enrichment analysis, and enrichment plot of one of the top-scoring hits is shown along with generative heatmap. B) Survival analysis (oncolnc.org) showing overall survival among available sarcoma cases within the TCGA database, low expression represents the lowest quartile, while highest expression represents a composite of the three higher quartiles, Log-rank P=0.05. C) ATAC-seq peaks were annotated according to genomic location (promotor transcription start site, transcription termination site, exon, intron, and intergenic) and plotted according to bright or dim aldeflour activity for the five sarcoma cell lines. D) Genome wide circos plot of ATAC-seq data generated from sorted soft-tissue sarcoma bright and dim cells. Increased ATAC-seq signal (accessible chromatin) directs outward, decreased chromatin signal directs inward. D) Increased chromatin accessibility at transcription start sites (TSS) correlating with the molecular target of EZH2. The plot on the right shows ATAC-seq signal as a ratio of bright to dim in WT GCT (UPS) cells was plotted for all canonical TSS in a 4 kb window, and ordered from low (top) to high (bottom). The plot on the left shows H3K27-me3 Chip-Seq data from a publicly available HepG2 dataset plotted according to the same rank-order as the right plot. Increasing black represents more Chip-Seq signal.

As an epigenetic modulator, increased EZH2 expression and activity leads to increased H3K27 methylation and subsequent gene silencing^16^. It has been demonstrated that EZH2 overexpression leads to suppression of differentiation or alternate cell fate pathways^17^. Consistent with this, it has been shown to maintain stem cell populations in both cancer and non-cancer tissues. EZH2 inhibition is an effective therapeutic strategy in epithelioid sarcomas that have loss of SMARCB1, a subunit of SWI/SNF complex that opposes the activity of EZH2^18^. Adding further support to the clinical relevance of EZH2, we evaluated EZH2 expression with respect to overall survival within The Cancer Genome Atlas (TCGA) dataset, which showed that the highest 75% of EZH2 expression was associated with significantly worse survival compared to sarcomas with the lowest quartile of EZH2 expression (Figure 3B, p=0.05)^19^. These data support further investigations of targeting EZH2 across sarcoma subtypes.

EZH2 modulates chromatin compaction through histone modification and therefore, we evaluated this in the cancer stem cell population using the Assay of Transposase Accessible Chromatin (ATAC-seq) which allows for an assessment of chromatin accessibility^16,20^. In parallel with Aldeflour-based sorting described above for RNA-Seq analysis, we also subjected the sorted material to ATAC-seq to evaluate differences in chromatin accessibility between bulk cells and CSCs. Following peak-calling and annotation, we noted a general trend in the cell lines tested for an increase in the number of total peaks, although on a percent basis there was little difference among different genetic regions between bright and dim cells (Figure 3C). This observation was further visualized using a circos plot of the fold differences between bright and dim cells representing CSCs and non-CSCs (Figure 3D). On a global level, and consistent with figure 3C, there was higher ATAC-Seq signal in bright cells compared to dim cells. The differences in chromatin accessibility were similar across sarcoma subtypes indicating a shared CSC specific program.

Having identified shared differential chromatin accessibility within CSCs, we next asked whether there was evidence for an EZH2-specific signature. To address this, we evaluated chromatin accessibility at transcription start sites in an unbiased manner across the genome and compared this with publicly available H3K27-me3 Chip-Seq data. We hypothesized that if EZH2 was involved in regulating chromatin accessibility in these areas, our ATAC-seq data would correlate with an increased H2K27-me3 chip-seq signal. We first ordered our gene list from a high to low ratio of ATAC-seq signal at TSSs, correlating to an ‘open’ vs ‘closed’ or ‘bound’ status, respectively. This ordered set of genes was then plotted against H3K27me3 signal from a publicly available HepG2 dataset, as a sarcoma specific dataset was not available. Consistent with our hypothesis, we observed increased H3K27-me3 chip-seq signal was increased in CSCs as compared to the non-CSC population (Figure 3E).

### Tazemetostat, a small molecule inhibitor of EZH2 abrogates the CSC phenotype

The EZH2 pathway is upregulated and there is an increase in H3K27-me3 across CSCs from different sarcoma subtypes. We then investigated if EZH2 was required to maintain the CSC phenotype. Interestingly, we were unable to generate viable EZH2-knockout cell lines using CRISPR, suggesting the importance of this pathway in sarcoma. Therefore, we utilized a small molecule inhibitor of EZH2 enzymatic activity, Tazemetostat, which is an S-adenosyl methionine (SAM) competitive inhibitor. Tazemetostat is FDA approved for the treatment of metastatic and locally advanced epithelioid sarcoma and is well-tolerated by patients. We first evaluated the effect of Tazemetostat in GCT cells representing undifferentiated pleomorphic sarcoma, the most common complex karyotype sarcoma in adults. At 48 and 72 hours, we observed little to no effect on overall cell viability (Figure 4A). The lack of short-term cytotoxicity in WT GCT cells was not unexpected, as we hypothesized that EZH2 specifically influences the CSC-phenotype. To investigate this possibility further, we performed soft-agar colony formation assays in cells treated with either vehicle or Tazemetostat. We found that continuous exposure to Tazemetostat significantly reduced the average size and abundance of soft-agar colonies (Figure 4B). These data suggested that inhibition of EZH2 by Tazemetostat blocks the CSC-phenotype in soft tissue sarcoma cells. To confirm this, we analyzed the CSC population using the Aldefluor assay. Extended treatment of GCT cells with Tazemetostat results in a decrease in the percentage of Aldefluor positive cells as compared to vehicle treatment. Removal of Tazemetostat showed that repression was transient and CSC populations rapidly returned to baseline. This data indicates that treatment with Tazemetostat specifically decreases the CSC population in soft tissue sarcomas (Figure 4C-D).

**Figure 4.**
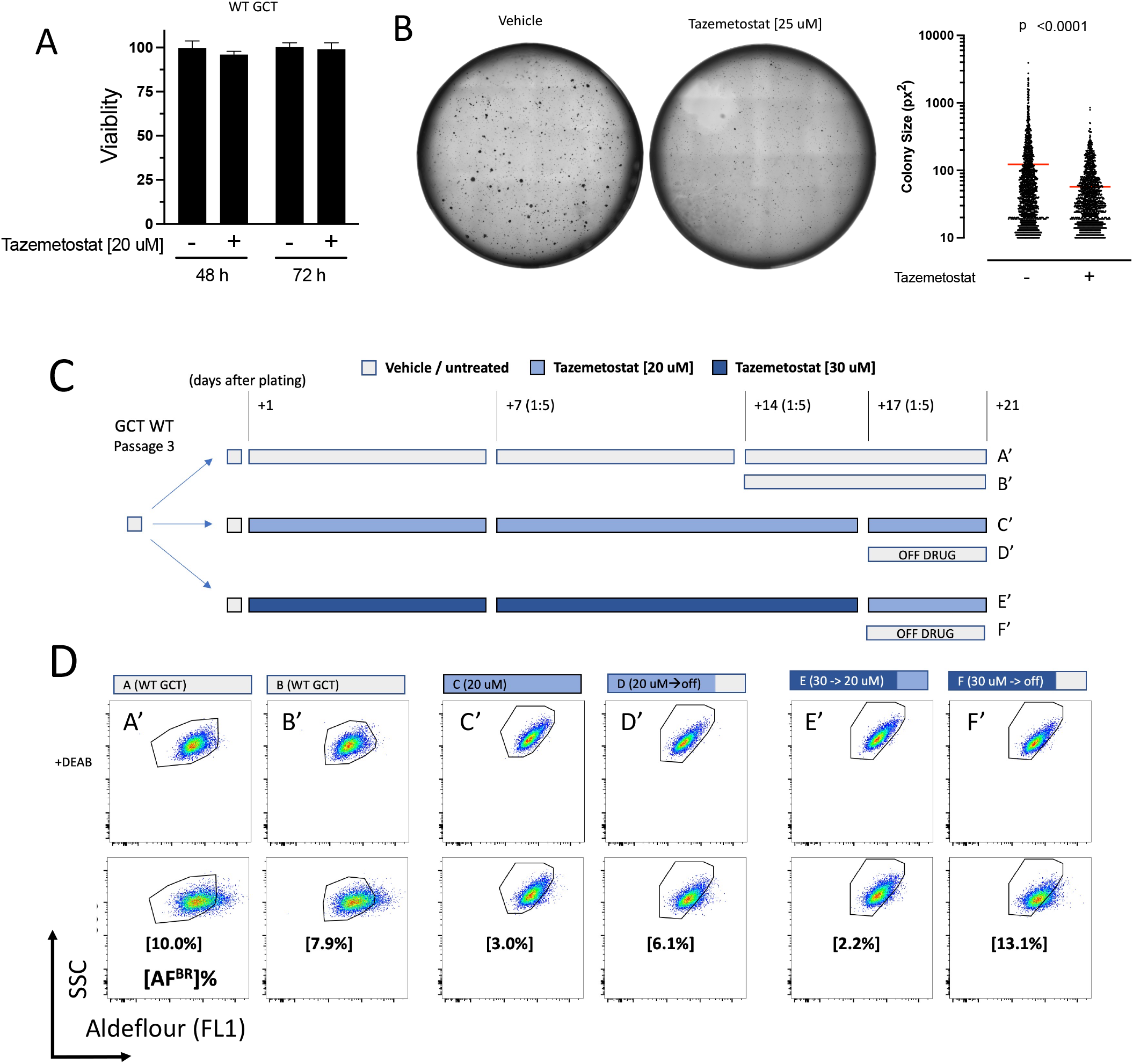
Inhibition of EZH2 with the small molecular inhibitor Tazemetostat abrogates the cancer stem cell phenotype. A) Tazemetostat does not significantly inhibit cell viability during short term treatment. Data are the mean±SD of three biological replicates, and are representative of three independent experiments. B) Representative soft-agar assay of WT GCT cells treated with vehicle or Tazemetostat (25 uM) (right) and quantification with imageJ (left). C) Treatment schema of WT GCT cells subjected to long-term culture with Tazemetostat. D) Aldeflour assay of WT GCT cells treated with vehicle or Tazemetostat over a 3 week period. DEAB controls for each experimental arm are shown in the top row, for which the corresponding gates are directly maintained in the bottom row. The percentage of aldeflour bright cells relative to DEAB controls is shown in brackets.

### Treatment with Tazemetostat sensitizes resistant cells to Doxorubicin

Given that resistance to doxorubicin was associated with an expansion of the CSC population and we found that inhibition of the epigenetic modulator EZH2 led to an ablation of the stem cell phenotype, we asked whether co-treatment of soft tissue sarcoma lines with doxorubicin and Tazemetostat would have a synergistic effect. To this end, we performed viability assays of WT and Doxo^R^ lines with doxorubicin and Tazemetostat alone or in combination (Figure 5). In both WT and Doxo^R^ SW872 cells, Tazemetostat had little appreciable effect on cell viability after treatment for 96 hours, consistent with earlier observations. While Tazemetostat in combination with Doxorubicin had a statistically significant synergetic effect in WT SW872 cells, this was much more pronounced in the Doxo^R^ derivative cells, completely rescuing the sensitivity of the cells to that of their WT counterparts (Figure 5A). We observed similar findings in WT and DoxoR GCT cells (Figure 5B). Taken together, these data support the hypothesis that EZH2 mediates the CSC phenotype in high-grade complex karyotype sarcomas and targeting of EZH2 sensitizes cells to anthracycline chemotherapy.

**Figure 5.**
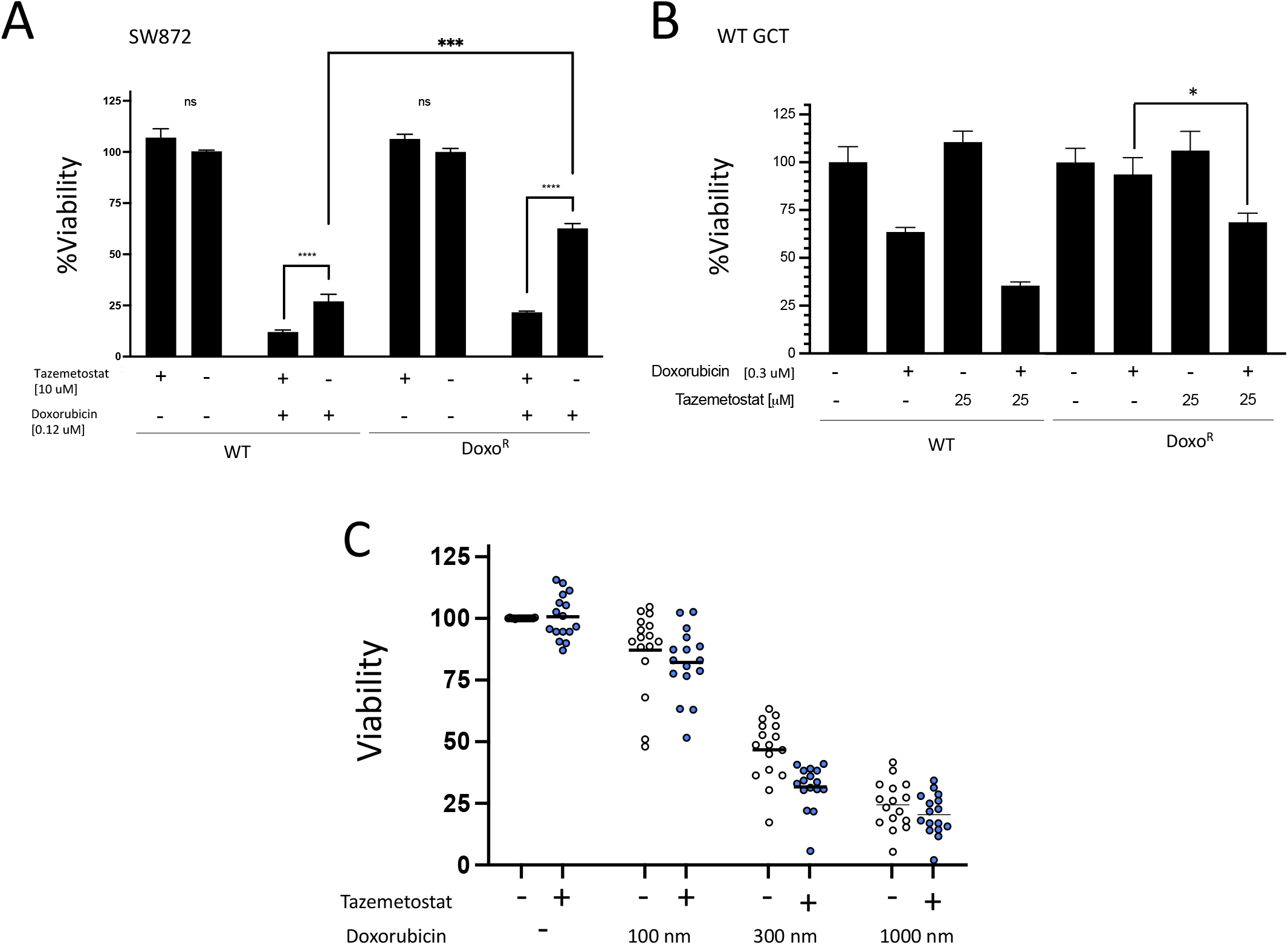
Co-treatment of soft tissue sarcoma cell lines with Tazemetostat and doxorubicin rescues treatment-acquired chemotherapeutic resistance. A) SW872 (LPS) and derivative doxo^R^ cells treated for 96 hours with doxorubicin and Tazemetostat alone or in combination. B) GCT (UPS) and derivative doxo^R^ cells treated for 96 hours with doxorubicin and Tazemetostat alone or in combination. Data for A-B are the mean ± SD of three biological replicates analyzed by ANOVA, * p<0.05, *** p<0.001 **** p<0.0001. C) Viability assays of unique, independently generated soft tissue sarcoma cell lines from oncogene-mediated forward transformation of mesenchymal stem cells *in vivo* treated with doxorubin at the indicated concentrations alone or in combination with 25 uM Tazemetostat. Data are the mean of triplicate biological replicates.

Having demonstrated a synergistic effect of doxorubicin and Tazemetostat in DoxoR lines, we wanted to determine if these results may be more broadly applicable. We recently developed an oncogene-drive forward genetics model in which human mesenchymal stem cells (MSCs) are transformed into high-grade soft tissue sarcomas^21^. From this, we have established a cell line panel of 14 independently derived, treatment-naïve sarcoma cell lines. These represent the undifferentiated pleomorphic sarcoma histologic subtype. We performed viability assays to determine whether these 14 unique and treatment-naïve sarcoma cell lines were uniformly or variably sensitive to doxorubicin, and whether co-treatment with Tazemetostat would potentiate these effects (Figure 5C). We appreciated variable sensitivity to doxorubicin among the cell lines, but noted that at all concentrations of doxorubicin tested (100, 300, and 1000 nM), the addition of 25 uM Tazemetostat potentiated the effects of doxorubicin at all concentrations tested for the vast majority of the sarcoma lines. Taken together, these data strongly support the synergistic effect doxorubicin and Tazemetostat in soft tissue sarcoma treatment, which can also be used to reverse acquired chemotherapeutic resistance.

## Discussion

In this study, we performed the first genetic and epigenetic analysis of CSCs across high-grade complex karyotype sarcoma subtypes to identify common therapeutic vulnerabilities. From this, we have identified EZH2 as a target to sensitize cells to anthracycline based chemotherapy. Tazemetostat, an EZH2 inhibitor is approved for treatment of SMARCB1 deficient epithelioid sarcomas^18^. This has been well tolerated by patients with a manageable side effect profile. The data provided here indicates that the use of Tazemetostat in conjunction with doxorubicin could be considered in the neoadjuvant or adjuvant setting to reduce the rate of recurrent disease. Additional approaches to overcoming drug resistance such as use of tyrosine kinase inhibitors or HDAC inhibitors have been shown to be efficacious and combinations of these therapies with Tazemetostat could be considered as a chemotherapy free approach^22,23^.

This study has demonstrated the importance of EZH2 in sarcoma CSCs and established EZH2 as a potential therapeutic target. Yet, the role of EZH2 in sarcoma CSC pathophysiology is not known and warrants further investigation. Sarcoma CSCs have active developmental pathways such as nanog, hedgehog, and hippo signaling. The regulatory overlap of EZH2 with these pathways could give additional insights into sarcoma CSC biology. In other tumor types, EZH2 has been shown to regulate key downstream targets to promote and expand the CSC population^17,24,25^. It is possible that the mechanisms by which EZH2 promotes the CSC population are similar but, given that EZH2 has specific cell type and context dependent functions, further investigation is needed.

One limitation of this study is that CSC markers have not been well characterized across sarcoma subtypes. Markers such as CD133 and CD117 have been identified in specific subtypes, but, expansion of these more broadly to additional subtypes has shown a lack of specificity for the stem cell population^26–29^. Additional subtype specific markers not been investigated across the disease spectrum. Therefore, in this study, we focused on Aldefluor positivity because this has been shown to identify a CSC population across tumor types, including multiple subtypes of sarcoma such as rhabdomyosarcoma and fibrosarcoma^30–32^. It is possible that Aldefluor may not identify all of the stem cells in each subtype. Further characterization of CSCs across this heterogeneous disease would allow for more detailed future studies.

Sarcomas are treatment resistant with high rates of recurrence and metastasis presumably due to the inability to eradicate the cancer stem population. This raises the possibility that targeting of this population could improve patient outcomes. In soft tissue sarcomas, the CSC population has not been well defined or extensively evaluated across sarcoma subtypes. The findings of this study create a foundation for further translational studies aimed at improving outcomes for patients with high-grade sarcomas.

## Materials and Methods

### Cell culture

SW872 cells (HTB-92) were purchased from American Type Culture Collection (ATCC, Manassas, VA) cultured in L-15 media supplemented with 10% FBS (Genesee Scientific, Morrisville, NC) in 0.1% CO_2_ atmosphere. GCT cells (TIB-223) were purchased from ATCC and grown in McCoys media supplemented with 10% FBS. All cells were cultured with antibiotic (pen/strep). SK-LMS-1 leiomyosarcoma cells were purchased from, ATCC (HTB-88). RH1 (755483-174-R-J1-PDC) non-alveolar rhabdomyosarcoma sample is a patient derived tumor cell culture from the NCI Patient-Derived Models Repository (PDMR, Bethesda, MD). Cells were cultured as described by the NIH protocol using DMEM/F12+Y compound. The myxofibrosarcoma sample is a PDX (918122-036-R) from PDMR. These were implanted into Non-SCID gamma (NSG) mice (Jackson Labs, Sacramento, CA) and once tumors reached 1mm they were harvested, digested in collagenase/dispase (Millipore Sigma, Burlington, MA) overnight to create a single cell suspension. Doxurubicin was purchased from Sigma and diluted in DMSO. Selection was performed by serial culture in concentration ranges from 0.03 uM initially, increasing by approximately 3 fold per selection step over the course of 3 months. Cells were passaged at dilutions of 1:5-1:10 every 2-3 days.

### Aldeflour assay

Cell culture populations representing cancer stem cells were identified using the Aldeflour assay (Stemcell Technologies, Cambridge, MA) according to the manufactures protocol, and optimized per cell line relative to reagent incubation duration. Assays were scaled using replicates as needed to achieve sufficient cell numbers for sorting.

### Flow Cytometry / Cell sorting

Flow cytometry was performed at the UC Davis Core Facility using BD Canto or Fortessa flow cytometers. Sorting was performed until sterile conditions using a BD Aria II sorter. Sorted cells were collected in respective serum-supplemented serum. Data was analyzed by FloJo software.

### Cell viability assays

Cell viability assays were performed using CellTiter Glo reagent (Promega, Madison, WI) according to the manufactuers recommended protocol. Cells were plated at a density of 2,000-10,000 cells per well in opaque 96-well white plates, cultured overnight, and then treated with serial dilutions of doxorubicin. Cells were then grown for the indicated time period, then assayed by Cell Glo-Titer. Luminescence was measured with a GloMax 96 luminometer (Promega).

### Soft agar colony formation assays

Soft agar colony formation assays were performed in 24 well plates. Stock solutions of 2x noble agar (1.2 and and 0.6 %) were prepared, sterilized, and stored at 4C until needed. On the day of preparation, stock solutions were warmed and the base layer was prepared by combining with a 2X solution of DMEM/F12, FBS, pen/strep (Genesee) and kept at 42C until used. The base layer (300 uL, 0.6% final agar concentration) was placed with pipette and allowed to solidy at room temperature. Cells were collected, counted, mixed as a 2X stock and combined with agar (0.3% final agar concentration, 10,000 cells/well), allowed to cool to room temp, and then covered with 100 uL of media. For quantification, wells were imaged in entirety using an EVOS M5000 microscope in a grid, stitched together using Image J software, thresholded, and then counted using the analyze particle function^33^.

### TCGA Analysis

Survival analysis linked to sarcoma-specific tumors was performed using an online web tool (Oncolnc.org)

### Xenografts

Xenograft tumors of SW872 and GCT cells (and their doxorubicin-resistant derivatives) were established in immunocompromised NSG mice (Jackson Labs Cat#005557) after injection of a 1:1 mixture of cells in PBS and Matrigel (1x10^6 cells/injection). All animal work was conducted at UC Davis and approved by the institutional animal care and use committee. Tumor growth was checked weekly and collected once palpable. Tumors were placed in formalin and processed for H&E staining by the pathology core at UC Davis.

### RNA Sequencing

Indexed, stranded mRNA-seq libraries were prepared from total RNA using the KAPA mRNA HyperPrep Kit (Roche) according to the manufacturer’s standard protocol for mRNA capture, fragmentation, random-primed first strand synthesis, second strand synthesis with dUTP marking, A-tailing, adaptor ligation, and library amplification. Libraries were pooled and multiplex sequenced on an Illumina NovaSeq 6000 System (150-bp, paired-end, >30 × 10^6 reads per sample). De-multiplexed raw sequence reads (FASTQ format) were mapped to the reference human transcriptome index (GRCh38/hg38, GENCODE release 36) and quantified with *Salmon*^34^. Gene-level read counts were then imported with *tximport*^35^. Differential expression analysis was performed with DESeq2^36^ and results were filtered for statistical significance (i.e., adjusted p < 0.05). Principal components analysis (PCA), hierarchical clustering, and Gene Set Enrichment Analysis (GSEA) (4,5) were performed on normalized read count data. For visualization of RNA-seq coverage tracks (bigWig) in the Integrative Genomics Viewer (IGV)^37^, read alignment was performed using the Spliced Transcripts Alignment to a Reference (STAR) ultrafast splice-aware aligner^38^ followed by generation of depth/control-normalized bigWig files with deepTools2^39^ using bins per million mapped reads (BPM) read coverage normalization.

### ATAC Sequencing

Sorted ALDH bright and dim cells (100,000) were used as input for library preparation. The protocol was followed to produce Illumina compatible libraries using the ATAC-seq kit from Active Motif. Briefly, cells were lysed and underwent tagmentation with Tn5 transposase and sequencing adapters. Illumina libraries sequenced using a NovaSeq 6000 (2 x 150bp, 45 x 10^6 reads per sample). Raw ATAC-seq data (FASTQ format) was analyzed using the nf-core/atac-seq pipeline (v1.2.1; https://nf-co.re/atacseq/1.2.1)^40^. This performed sequence read quality control analyses with FastQC^41^, adapter trimming, mapping to the reference human genome assembly (GRCh38/hg38) with Burrows-Wheeler alignment^42^, and post-alignment data cleanup steps, such as duplicate marking (picard) and read filtering. Normalized bigWig files were created and passed to deepTools^39^ to generate gene-body metaprofiles and calculate genome-wide enrichment. Peaks calling was performed with the Model-based Analysis of ChIP-Seq (MACS2) algorithm^43^ followed by annotation with HOMER (Hypergeometric Optimization of Motif EnRichment)^44^, and creation of consensus peaksets from across all samples with BEDTools utilities.

## Data Availability

RNA sequencing and ATAC-seq data are publicly available at the Gene Expression Omnibus repository (pending).

## Author Contributions

EO was responsible for the overall scientific question and experimental design, performing experiments, data analysis, writing of the initial draft. MM performed experiments, extracted and analyzed data. CT and RD were responsible for analysis and interpretation of bioinformatics experiments. JCA was responsible for the overall scientific question and experimental design, obtaining funding, writing the manuscript.

## Acknowledgements

JCA is funded by the National Cancer Institute/National Institutes of Health grants 5K12-CA138464 and National Cancer Institute/National Institutes of Health grant P30CA093373, in addition to the Doris Duke Charitable Foundation/Burroughs Wellcome Fund and Landgraf foundation. Genomics support (CT and RD) and Flow Cytometry faciltiies were funded by the UC Davis Comprehensive Cancer Center Support Grant (CCSG) awarded by the National Cancer Institute (NCI P30CA093373). The authors thank Qian Chen from the UC Davis Center for Genomic Pathology for histology support.

